# Middle-infrared Modulation Of Sleep By Cardiometabolic Function

**DOI:** 10.1101/2023.11.28.569000

**Authors:** Yan Sun, Guang Xian, Chao Wang, Delong Zhang

## Abstract

The interdependence of sleep and overall well-being is profound. Sleep has the capacity to regulate hematopoiesis, thereby sustaining our physiological equilibrium. Nevertheless, our understanding regarding the potential reversibility of this regulatory process remains limited. In this study, we present evidence suggesting that exposure to middle-infrared radiation (MIR) within the 5-7 mm wavelength range can elicit non-thermal biological effects within the human body. These effects are characterized by an augmentation in blood circulation, mediated through modifications in red blood cell properties. Furthermore, MIR exposure is accompanied by a reduction in homocysteine levels and suppressor T-cell populations. Importantly, daytime modulation of MIR has been shown to significantly facilitate the amelioration of insomnia symptoms during nighttime, primarily through its impact on the architecture of deep sleep. Conclusively, our investigation reveals a robust association between cardiometabolic function and human sleep, with MIR presenting a promising avenue for enhancing sleep quality by regulating both the circulatory system and the immune system. These findings pave the way for the development of therapeutic strategies aimed at mitigating prevalent sleep-related disorders by disentangling the intricate links between sleep and cardiometabolic function.

## Introduction

Sleep is a vital physiological and psychological requirement for maintaining a balanced state in both body and mind. It plays a crucial role in the overall well-being of an organism. (Tufik, Andersen, Bittencourt, & Mello, 2009) Insufficient sleep has been found to elevate the chances of developing cardiovascular diseases by affecting the immune system. (Cappuccio, Cooper, D’elia, Strazzullo, & Miller, 2011) Sleep also has the ability to regulate hematopoiesis and provide protection against atherosclerosis. (McAlpine et al., 2019) Despite recognizing the strong connection between sleep and cardiometabolic function, it remains largely unclear whether modulating cardiometabolic activity can effectively improve sleep quality.

All living organisms on Earth benefit from the infrared radiation emitted by the Sun, as it promotes biological effects. The red blood cell (RBC) is the specific target of this infrared radiation action.(Chludzińska, Ananicz, Jaros ławska, & Komorowska, 2005; Iijima, Shimoyama, Shimoyama, & Mizuguchi, 1991; Piasecka, Leyko, Krajewska, & Bryszewska, 2000; Sadowska, Krajewska, & Bryszewska, 2001; Tomasz Walski, Chludzi ń ska, Komorowska, & Witkiewicz, 2014) The investigation of the effects of photobiomodulation on RBC properties can contribute to a deeper understanding of the interconnected regulation mechanisms between sleep and cardiometabolic function. Numerous studies have shown that the utilization of infrared radiation (with a wavelength range between 4-16 µm, as defined by ISO 20473) plays a significant role in promoting various biological effects at different levels, ranging from sub-cellular organelles to human behavior in a diverse range of species.(Vatansever & Hamblin, 2012) The exposure of RBC to infrared radiation can induce changes in cellular properties by impacting membrane fluidity and membrane potential. This may potentially contribute to the improvement of microcirculation and metabolism in the human body. (T Walski et al., 2015) Until recently, the investigation of cellular properties influenced by infrared radiation has been the primary focus. Considering the absorption spectra of water, it seems reasonable to consider water as the most suitable candidate. (Burns & Ciurczak, 1992; Libnau, Kvalheim, Christy, & Toft, 1994) Additionally, the structural changes in water bound to the cell surface caused by dehydration can result in alterations in membrane protein conformations. (Knight, Goodall, & Greenhow, 1979; Natzle & Moore, 1985; Phillips & Eyring, 1986) The impact of infrared radiation on human erythrocytes plays a key role in promoting accelerated microcirculation and enhancing cardiometabolic function for biological effects. (Iijima et al., 1991; Komorowska, Cuissot, Czarnoleski, & Bialas, 2001) Despite the widespread use of infrared radiation therapy in various clinical applications, such as wound healing(Whelan et al., 2001) and treatment of neurodegenerative disorders(Johnstone, Moro, Stone, Benabid, & Mitrofanis, 2016), researchers universally acknowledge that the therapeutic benefits strictly rely on the dosage of infrared radiation received by the cells.

In this study, we report that the mid-infrared radiation (MIR) with a wavelength of 5-7 mm has the ability to induce significant non-thermal biological effects, resulting in changes to red blood cell (RBC) properties at room temperature during the daytime. Furthermore, we demonstrated that MIR therapy can effectively alter the sleep architecture in patients with chronic insomnia, aiding in the recovery of insomnia symptoms during the night. These findings serve to complement recent research that has shown how mid-infrared stimulation (wavelength of 5.6 mm) can exert reversible and non-thermal modulatory effects on ion channel activity, neuronal signaling, and sensorimotor behavior. Such findings suggest that mid-infrared neuromodulation may hold promise as a therapeutic approach. (Liu, Qiao, Chai, Zhu, & Shu, 2021) In addition, this study provides further evidence supporting the use of photobiostimulation targeting RBCs as a safe, effective, and widely applicable method for generating therapeutic effects. Through modulation of sleep by cardiometabolic function, mid-infrared radiation offers a potential avenue for intervention in sleep disorders. Overall, this study highlights the substantial impact that MIR can have on biological systems and its potential applications in clinical settings.

## Results

### Non-thermal biological effect of MIR

Numerous studies have revealed that exposure of RBCs to infrared radiation can result in alterations of cellular membrane fluidity and membrane potential in living organisms.(T Walski et al., 2015) Based on the previous research findings, we investigated the biological impact of MIR on the human body. Initially, we examined the pattern of RBC distribution before and after exposure to MIR in healthy participants using a microscope equipped with a photographic device. Our analysis revealed a significant reduction in RBC aggregation following a 20-minute MIR session (Figure 1A, Figure S1), and even a 5-minute MIR session also showed similar effects (Figure 1B, Figure S2). This change in RBC aggregation is likely attributed to alterations in RBC properties caused by direct blood photobiomodulation from low-level light therapy in the MIR spectral region, resulting in modifications to membrane fluidity and membrane potential. (Chludzińska et al., 2005; Komorowska et al., 2001; T Walski et al., 2015)

**Figure 1.**
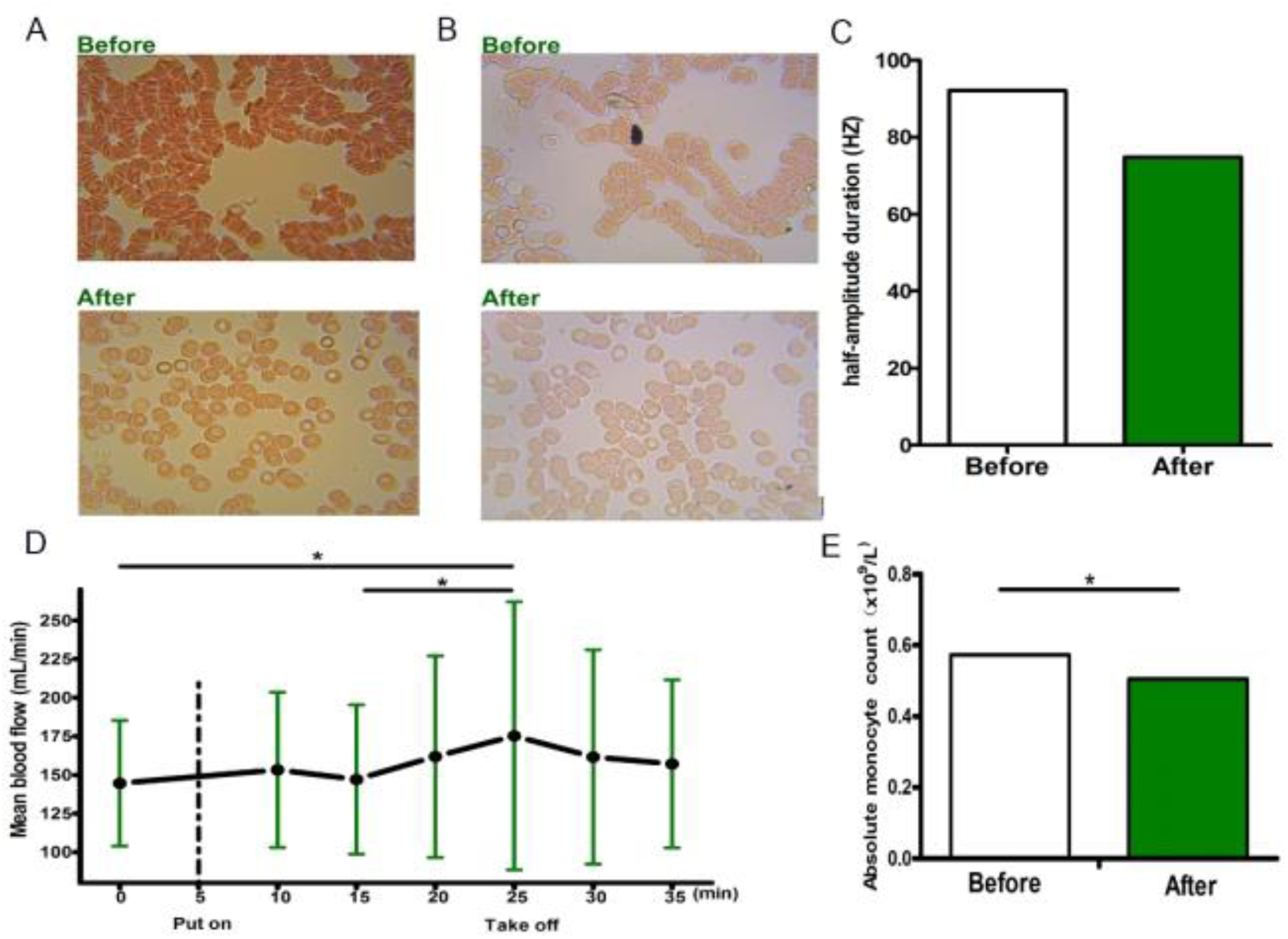
Non-thermal biological effect of MIR. (A) RBC properties change from 20 minutes bracelet wearing; (B) RBC properties change from 5 minutes bracelet wearing; (C) Half-amplitude duration of water changes from 20 minutes bracelet wearing; (D) Blood flow dynamic changes before and after 20 minutes bracelet wearing; (E) Absolute monocyte count changes from 20 minutes bracelet wearing. \**P* < 0.05, \*\**P* < 0.01, \*\*\**P* < 0.001.

In addition to assessing RBC aggregability, we took our investigation a step further by examining blood flow using laser speckle contrast imaging in the MIR region. The findings revealed that MIR stimulation significantly enhanced blood flow, with the highest increase observed after 20 minutes of accumulation (*P* = 0.036, as determined by a one-tailed paired *t*-test; see Figure 1D). These results align with previous research demonstrating the influence of RBC aggregation on blood flow properties.(Baskurt & Meiselman, 2003; Keymel, Heiss, Kleinbongard, Kelm, & Lauer, 2011; Lowe, 1987) However, when considering the collective data, it becomes evident that the use of MIR has the capability to induce a noteworthy impact on the cardiometabolic function in the human body.

### MIR-derived blood quality change

*Immediate effect*. In order to explore the MIR-derived blood quality change from the short-term action, we compared the results of a routine blood test before and after 20-minutes MIR on 15 healthy participants, and we found that the absolute monocyte count was significantly changed (*t* = -2.743, *P* = 0.013, Figure 1E).

#### Chronic effect

In order to track the follow-up effect of the MIR-derived biological effect on human body, we explored the modulation effect of chronic MIR on the blood properties in the patients with renal failure. We compared the results of routine blood test before and after 4-week MIR (MIR lasting for half an hour in the morning, noon, and evening), and did a 2-week follow up study to monitor routine blood test, i.e., Homocysteine (HCY) and T-cells. We found that the MIR significantly increased Mean Cell Hemoglobin Concentration (MCHC) (M*diff* = 51.111, *P* = 0.003), Mean Corpuscular Hemoglobin (MCH) (M*diff* = 3.356, *P* = 0.054), Basophil (BA) (M*diff* = 0.019, *P* = 0.014) and Basophil percent (BA%) (M*diff* = 0.367, *P* = 0.028), and significantly reduced Red blood cell distribution width (RDW) (M*diff* = -5.344, *P* = 0.011) at the fourth week. In addition, we found that the MIR significantly increased MCHC (M*diff* = 50.111, *P* = 0.002), MCH (M*diff* = 3.189, *P* = 0.045), BA (M*diff* = 0.029, *P* = 0.041) and BA% (M*diff* = 0.456, *P* = 0.022), and significantly reduced RDW (M*diff* = -4.533, *P* = 0.017) at the sixth week. Of note, the promotion effect of MCHC, MCH, RDW and BA% were stable between the fourth and the sixth week. Importantly, the results showed that the MIR significantly decreased the suppressor T-cells percentage (26.94 ± 8.47 vs 25.746 ± 8.75, *t* = 2.794, *P* = 0.021), CD3+CD8+%Lymph (26.94 ± 8.47 vs 25.728 ± 8.76, *t* = 2.853, *P* = 0.019) and the HCY (12.75 ± 2.981 vs 9.86 ± 2.949, *t* = 2.947, *P* = 0.016) (Figure 2).

**Figure 2.**
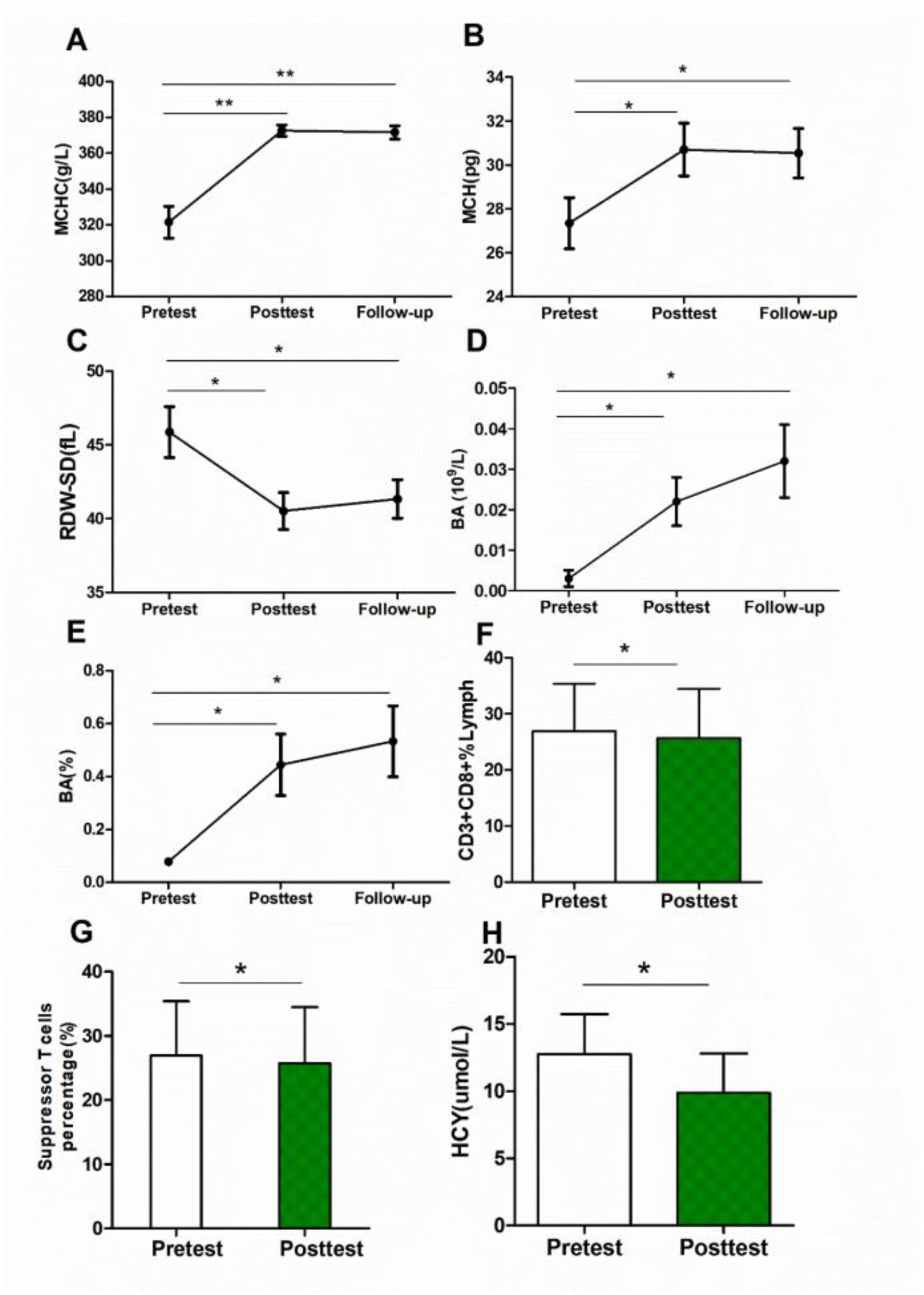
MIR derived blood quality changes from 4-week and/or follow-up 2-week bracelet wearing. (A) MCHC; (B) MCH; (C) RDW; (D) BA; (E) BA%; (F) CD3+CD8+%Lymph; (G) Suppressor T-cells percentage; (H) HCY. \**P* < 0.05, \*\**P* < 0.01, \*\*\**P* < 0.001.

The change of the RBC properties accompanying with the immune cell and the platelet change may result in the improved oxygen delivery ability during blood circulation, which was corresponding to many previous studies that the RBC aggregation would impact blood viscosity, blood flow resistance/velocity,(Yalcin, Meiselman, Armstrong, & Baskurt, 2005) and even the functional capillary density,(Vicaut, Hou, Decuypère, Taccoen, & Duvelleroy, 1994) all of which influence the blood flow oxygen delivery-especially ability in micro-circulation.(Barshtein, Benami, & Yedgar, 2014)

### MIR biological effect modified human sleep anatomy

#### Sleep is sensitively regulated by the MIR

We further explored the MIR biological effect on the insomnia symptom recovery. By using the Pittsburgh Sleep Quality Index (PSQI) test, we measured the change of the insomnia symptoms flowing with 4-week MIR. We found that the PSQI scores were significantly decreased in the experimental group (10.55 ± 3.02 vs 4.95 ± 2.39, *t* = 10.870, *P* < 0.001), which was not observed in the control group (13.05 ± 4.16 vs 12.9 ± 4.08, *t* = 0.719, *P* = 0.481, Figure 3A). By combining the PSQI and the Polysomnography (PSG) to explore the sleep anatomy change underlying the insomnia symptom recovery from 4-week MIR. We found that the MIR significantly advanced sleep quality on the PSQI scores (Global score: 9.48 ± 2.68 vs 2.97 ± 1.90, *t* = 17.003, *P* < 0.001, Figure 3C).

**Figure 3.**
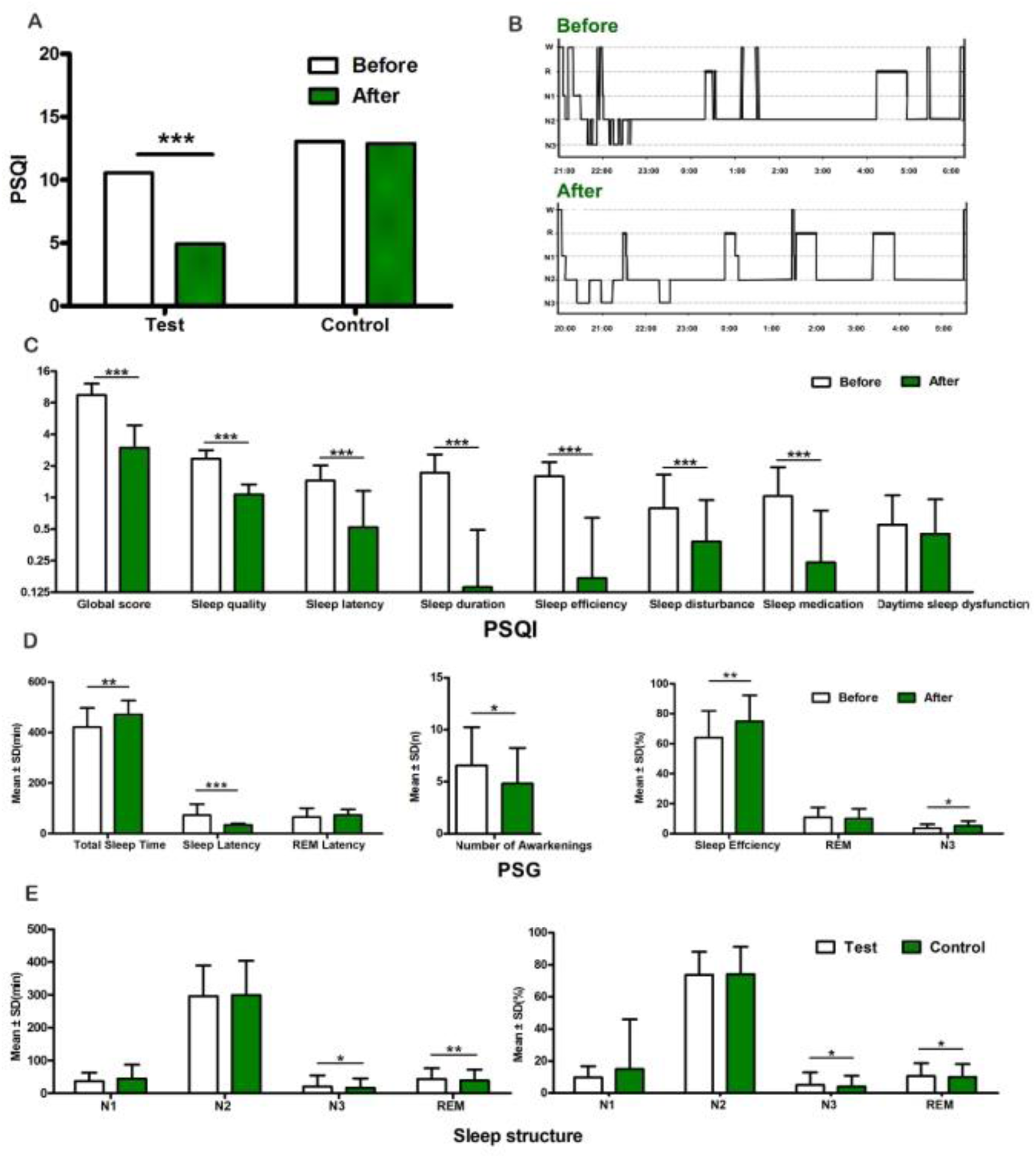
Modified human sleep anatomy from MIR with and without MIR. (A) PSQI scores; (B) Sleep anatomy; (C) PSQI results; (D) PSG results; (E) Sleep anatomy. \**P* < 0.05, \*\**P* < 0.01, \*\*\**P* < 0.001.

The PSG showed that the MIR significantly decreased the sleep latency (72.56 ± 43.56 vs 33.62 ± 5.04, *t* = -5.789, *P* < 0.001) and the number of awakenings (6.57 ± 3.67 vs 4.81 ± 3.43, *t* = -2.219, *P* = 0.032), giving rise to the sleep efficiency (64.07 ± 17.71 vs 74.91 ± 17.34, *t* = 3.193, *P* = 0.003) and the total sleep time (420.77 ± 76.06, 471.25 ± 55.00, *t* = 3.167, *P* =0.003, Figure 3D). Further analysis showed that the MIR obviously increased the percentage of the N3 stage (3.58 ± 2.69 vs 5.20 ± 3.16, t = 2.561, *P* = 0.014, Figure 3, B and D) of the entire sleep process. And we measured the change of the PSG flowing with 4-week comparing test (MIR lasting for half an hour in the morning, noon, and evening) and control (fake MIR lasting for half an hour in the morning, noon, and evening) groups. And we found that the MIR significantly increased the percentage (4.00 ± 6.78 vs 5.13 ± 7.72, *t* = 1.136, *P* = 0.027) and the time of N3 stage (16.61 ± 28.95 vs 20.81 ± 33.33, *t* = 0.961, *P* = 0.038), the percentage (9.99 ± 8.00 vs 10.62 ± 7.86, *t* = 0.188, *P* = 0.021) and the time of REM stage (38.51 ± 33.41 vs 43.26 ± 32.79, *t* = 1.026, *P* = 0.006) (Figure 3E).

Finally, we recruited another 350 chronic insomnia patients (age range: 24-83 years, 168 males, mean insomnia history is over 1 month) to trace the dynamic change of the improvement of sleep symptoms coming from the MIR biological effect. The MIR setting were corresponding to thatas aforesaid, and the self-report (1-5 scale representing no symptom to very severe) was used to measure the symptom change. The results indicated that the significant insomnia symptom recovery was observed at about the 2th week. At the 4th week, the ratio of the symptom recovery reached 59.5%, and the ratio of mild symptoms reached 33.5% (Figure 4). The observation showed that MIR during the daytime facilitates a good quality sleep at night.

**Figure 4.**
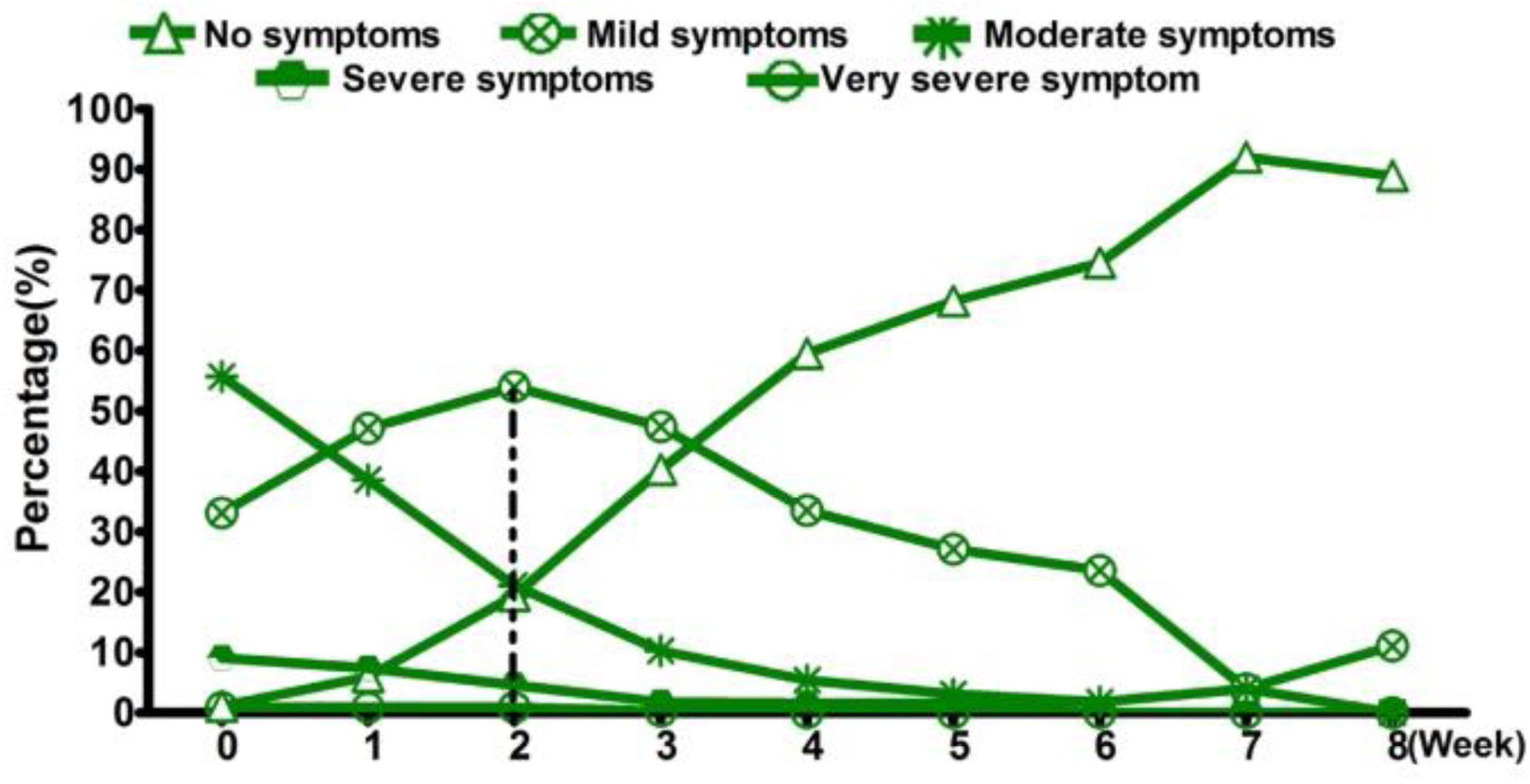
Insomnia symptom recovery from MIR.

## Discussion

The current study, to the best of our knowledge, presents the pioneering neurophysiological mechanism and behavior metric providing experimental evidence that the MIR biological effect effectively modifies cardiometabolic function, facilitating human sleep behavior. Despite numerous studies reporting the beneficial effects of infrared radiation on the human organism, little has been known about its biological modulation effect on the blood circulation system and its impact on human sleep. Consequently, the results demonstrate the initial evidence of the significant improvement in the dose of infrared radiation received by cells due to the MIR effect. Moreover, we also highlight one crucial pathway through which the infrared radiation effect targets RBC and alters micro-circulation, aiming to rehabilitate human sleep symptoms. Prior research has demonstrated the modulatory effect of sleep on hematopoiesis. The current data not only support previous findings but also enhance our understanding of the bidirectional modulation between sleep and cardiometabolic function.

There is a neuro-immune axis directly connects sleep to immune function and cardiovascular disease, and the undisturbed sleep modulates haematopoiesis to protect against atherosclerosis(McAlpine et al., 2019). To uncover the essence of the tight relation between sleep and blood circle is critical since it is related to the essential physiological metabolism of life, in which to determine the research target is the key point. On the one hand, water appears to be the most rational target as it is omnipresent in cells. With the knowledge growth of aquaporins on cell membranes, researchers have realized that the absorption of water by cell membranes has a certain selectivity, which not only depends on traditional permeation, but also is closely related to the proteins on cell membranes. Thus, changing the properties of water to modulate cell properties has become an important research hotspot. On other hand, RBC is the most abundant type of blood cells in the human body and obtains a long circulation time (120 days in human),(Barshtein et al., 2014; Oldenborg et al., 2000) and RBC is not only the inert oxygen carriers, but also is acting as an important modulator of the innate immune response.(Darbonne et al., 1991; Hotz et al., 2018; Neote, Darbonne, Ogez, Horuk, & Schall, 1993) Of importance, MIR effect has been demonstrated to modulate RBC properties.

With the advantage of the MIR effect, we could optimize the blood quality with a high efficiency to explore the modulation effect from blood to sleep. As we noted that the mice with abnormal sleep could produce more monocytes to develop larger atherosclerotic lesions.(McAlpine et al., 2019) In the present study, we firstly showed that the MIR could change RBC properties and blood circle performance, and also could decrease absolute monocyte count and other cells in immune systems. There were many researches focusing on the RBCs as important modulators of the innate immune response. (Buttari, Profumo, & Riganò, 2015) The present findings might show that there may exist a bidirectional relation between sleep and blood circle in which the immune system may take an important role in maintaining the balance of them. To confirm the key role of immune system in infrared modulation of sleep by cardiometabolic function, we further observed that the HCY and the T cell amount were significantly changed before and after 2-week bracelet wearing. Until recently, HCY may contribute to the development of atherosclerosis through increasing endothelial cell interaction with monocytes and T cells by increasing the expression of adhesion molecules.(Koga, Claycombe, & Meydani, 2002) Of note, the hyperhomocysteinemia has been observed to contribute to cardiovascular diseases in the presence of other risk factors such as inflammation, a known risk factor for cardiovascular diseases, that the elevated levels of HCY contribute to inflammation, in which the HCY, T cell immunity, and hypertension act together in increasing the risk for cardiovascular diseases.(Veeranki, Gandhapudi, & Tyagi, 2016) The observed reduced HCY and the inflammation reaction might show the action mechanism of MIR on the cardiometabolic function.

By using the PSG, we found that MIR significant induced the increase of the Non-rapid eye movement sleep (NREM), especially the deep sleep (N3). As we know, the NREM state related to the slow-wave activity (SWA, 0.5–4.5 Hz), to the extent, reflects the homeostatic regulation of sleep need, the SWA was usually used as an index of synaptic homeostasis,(Vyazovskiy, Cirelli, Pfister-Genskow, Faraguna, & Tononi, 2008) during sleep, which is both a sensor and an effector in a homeostatic process occurring.(Tononi & Cirelli, 2014) Corresponding to the previous findings of biochemical indicators, we further found that the MIR effect significantly reduced the HCY and the inflammation reaction of the immune system to decrease the stress response of the human body at the daytime, which could reflect on the sleep anatomy at night. A recent study has shown that the regular sleep could modulate haematopoiesis by the way of immune system,(McAlpine et al., 2019) in response to this study, we suggests that the MIR regulation of hematopoietic function can in turn improve sleep, which was also through the immune system pathway.

In the present study, we provided experimental evidence for that the MIR effect could effectively change the cardiometabolic function. Meanwhile, because this intervention works by regulating the blood circulation system and improving the immune system to eliminate stress response, thus, a short-term MIR during the daytime can improve sleep at night, which means MIR during the daytime could facillate a good sleep at night.

Our data thus establish a tight relation of cardiometabolic function to human sleep behavior, and that the MIR could profit human sleep by regulating the blood circulation. Given the existence of a vicious circle of sleep and cardiovascular problems, these results identify a safe strategy to reverse the links between sleep and cardiometabolic function, and may lead to new therapeutic strategies for treating the related highly prevalent disorders.

### Limitations and future directions

There were several limitations that should be addressed in future research. Firstly, this study validated the role of MIR modulation in enhancing sleep through improved blood circulation, and proposed a potential biochemical mechanism. However, additional investigations are needed to elucidate the entire chain of biochemical reactions, particularly with regards to ion channels, in order to provide more detailed evidence. Secondly, sleep behavior has been shown to be closely linked to neural activity in the brain. Therefore, future studies should employ EEG and fMRI technology to investigate the underlying neural mechanisms. Lastly, the objective of this study was to enhance blood circulation by increasing the absorptivity of red blood cells to far-infrared rays, thereby improving sleep quality. Hence, further research is required to determine if this mechanism can be applied to the treatment of other diseases.

## Methods and materials

### Materials

The MIR-emitted material was extracted from the calcination of rare earth of the lanthanide series, nanometer titanium dioxide, and zinc dioxide under a high temperature condition (1500 ℃) flowing with 12.5-25 MPA in the high-temperature and pressure autoclave lasting for 43 hours (Fig. 5A). The MIR emitted by this material has a maximum emission wavelength at about 8 um (20 ℃, 41 RH) (Fig. 5F). The 6-Hz audible sound beats on 1,000 Hz carrier tone was applied to promote the MIR biological effect (Fig. 5C). We put the MIR-emitted material and the audible sound beat setting together as a facility (Fig. 5E) which was worn at the medial wrist of the left hand (about three fingers away from the palm, Fig. 5B) to deliver the amplified MIR (Invention Patent Number Authorized by Chinese Government : ZL 2020 1 1604234.9).

**Figure 5.**
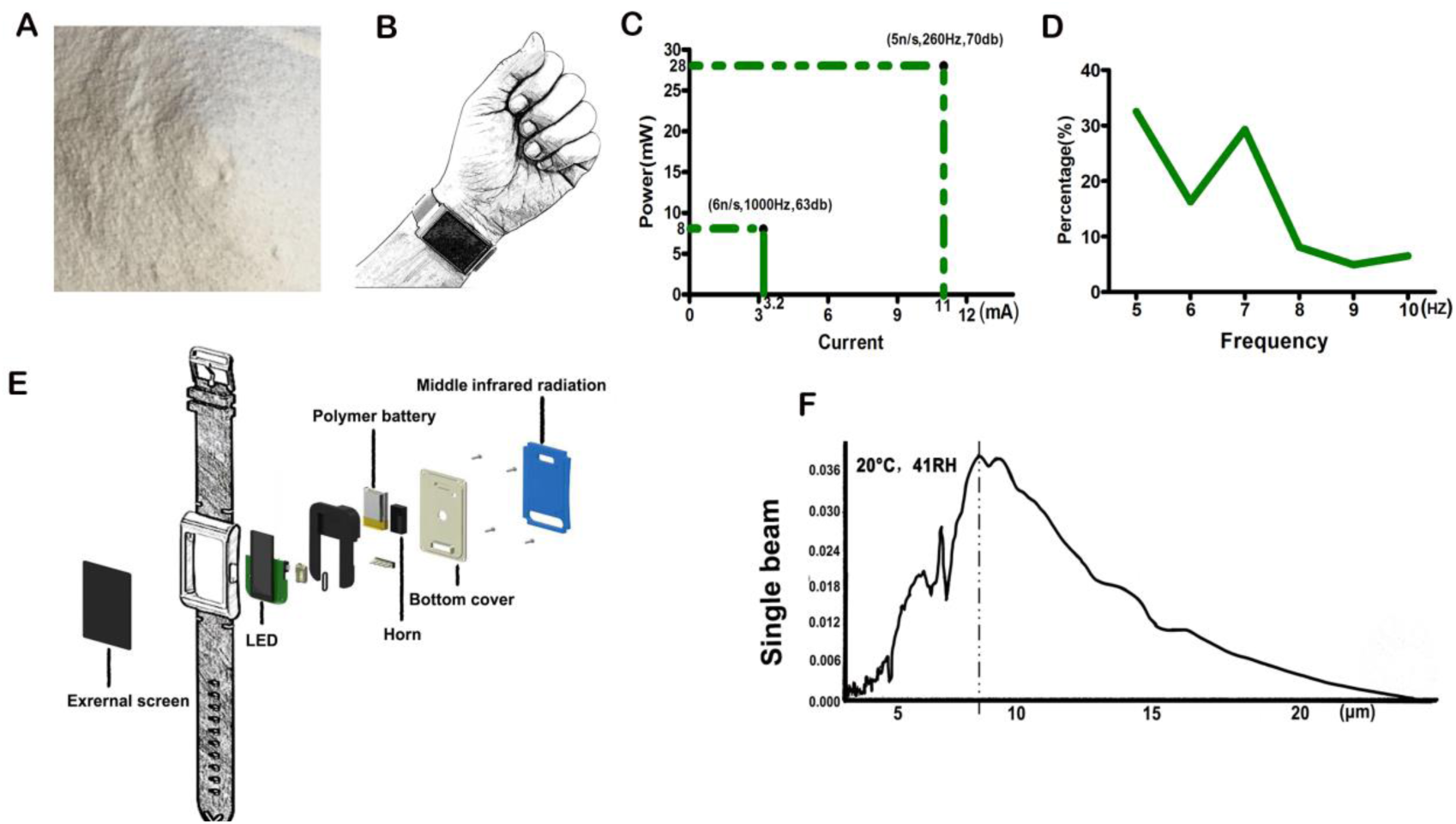
Materials and setting. (A) The extracted MIR-emitted material; (B) Manner of bracelet wearing; (C) Audible sound beat parameter setting; (D) The percentage of the selected sound beats frequency; (E) The bracelet combining the MIR material and the audible sound beat laugher; (F) MIR wavelength.

A type of hot-pressed polycrystalline semiconductor material, characterized by its raw materials consisting of, by weight, inorganic powder 10-33 parts, silicate 5-15 parts, rare earth materials 2-10 parts, polymers 15-30 parts, permeating agents 5-15 parts, and solvents 35-65 parts. The said inorganic powder comprises inorganic oxides; the inorganic oxides are selected from titanium dioxide, zinc oxide, silicon dioxide, and aluminum oxide. The material preparation includes the following steps:

1. The inorganic powder, rare earth materials, and silicate are separately pulverized, mixed uniformly, and calcined in a high-temperature furnace at 1200°C for 60-80 minutes to obtain mixture 1;
2. Mixture 1 is added to a reaction kettle to react for 16-20 hours, where the temperature is 900-1000°C and the pressure is 10-15 MPa, then cooled and ground to obtain mixture 2;
3. Mixture 2 prepared in step (2), and the permeating agent are stirred in the solvent for 2-4 hours to obtain mixture 3, then the elastomer is heated to 160-170°C, mixture 3 is added, stirred for 10-15 hours, and granulated in an extruder to obtain the final product. By setting the sintering temperature, reaction temperature, and reaction pressure reasonably, the hot-pressed polycrystalline semiconductor material prepared by the method in this technical solution has a high yield, good mechanical strength, and is safe, free from free radiation, and heavy metal radiation; its emitted far-infrared wavelength range is 5-7μm, with a peak at 6μm. It provides a kind of silicon-based semiconductor material that can generate 5-7μm (with a peak at 6μm) under normal temperature with the acousto-optic effect, as shown in (Fig. 6A), and has a redshift effect. It lays the foundation for smart wearable health functionalities and opens a new path for treating diseases through physical methods via an oxygen solubility channel.

**Figure 6.**
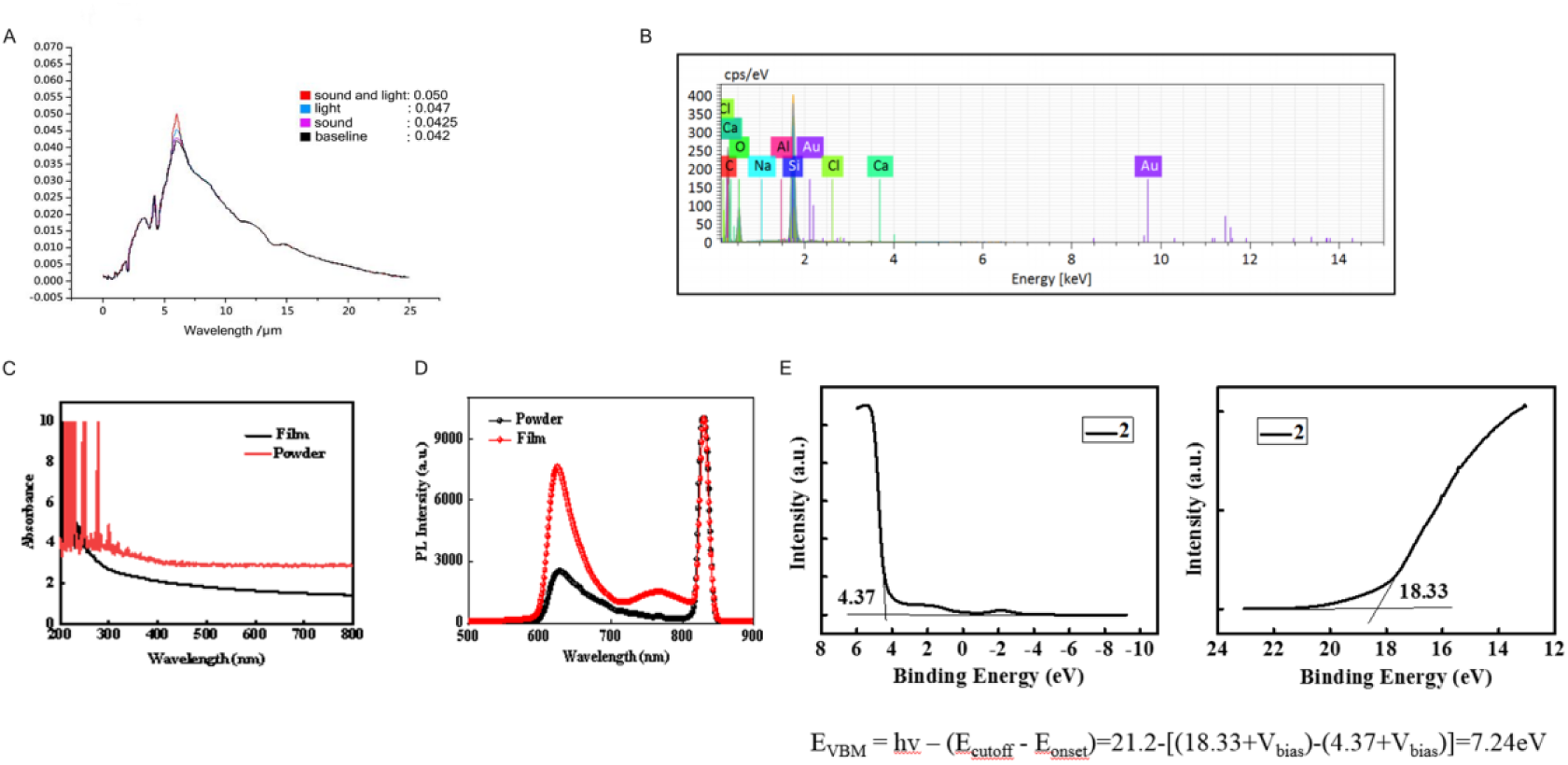
Material preparation. (A) Under normal temperature with the acousto-optic effect; (B) Contents of a novel semiconductor material; (C) Toptical properties of the thin films; (D) Toptical properties of the target compound powder; (E) Ultraviolet Photoelectron Spectroscopy (UPS) test of the thin films

In this project, through a series of high-temperature and high-pressure reactions, the person in charge obtained a novel semiconductor material. SEM-EDS analysis revealed that the target compound contains elements such as C, O, Na, Al, Si, Cl, Ca, etc., with contents as shown in (Fig. 6B). Systematic characterization of the optical properties of the target compound powder and thin films was conducted through UV-Visible spectroscopy and fluorescence spectroscopy. As shown in (Fig. 6C), the thin-film samples exhibit a weak absorption peak in the UV region, with the calculated optical bandgap being 3.72 eV, indicating a direct bandgap semiconductor. Meanwhile, fluorescence spectroscopy showed a clear fluorescence emission peak for the powder samples between 600 to 800 nm (Fig. 6D), with the absorption peak at 626 nm, corresponding to an energy of 1.98 eV. Corresponding thin-film samples exhibited two distinct absorption peaks between 600 to 700 nm and 700 to 800 nm, with peak values at 624 nm (1.99 eV) and 767 nm (1.62 eV). The valence band top was found to be at 7.24 eV, as analyzed and calculated from the Ultraviolet Photoelectron Spectroscopy (UPS) test of the thin films (Fig. 6E).

### Participants

The present study recruited a total of 17 healthy participants, 12 with chronic renal failure patients with stage 5 chronic kidney disease at the Yong’an hospital in Foshan, and 590 chronic insomnia patients at Henan Mental Hospital in Xinxiang.

These healthy participants did not have any history of cognitive disorders, neuropsychiatric disorders, or drug abuse. We examined the blood flow dynamic change induced by the MIR on 17 healthy participants (age range: 25-61 years, 6 male) by using the laser speckle contrast imaging (LSCI).

The patients with chronic renal failure neither had systemic anticoagulation contraindications nor had a suitable family environment for hemodialysis support. The patients with pregnancy, lupus erythematous, rheumatoid arthritis, or those need to prevent antibiotics were excluded s for this study. We recruited 12 patients with chronic renal failure (age range: 32-65 years, 9 females), and compared the results of routine blood test before and after 4-week wearing, and then did a 2-week follow-up study to monitor the routine blood test (i.e., HCY and T-cells).

In total of 590 chronic insomnia patients (aged from 18 to 85 years) were collected according to DSM-IV criteria, including: 1) Sleep disorder is almost the only symptom, and other symptoms are secondary to insomnia, including difficulty in falling asleep, insomnia-middle, easy to wake up, more dreams, not easy to fall asleep again after waking up, feeling uncomfortable after wake up, fatigue or daytime sleepiness and so on; 2) These symptoms of sleep occurred at least three nights a week for at least 3 months and insomnia causes significant severe distress or discomfort or impedes social function; 3) Of note, the participants who meet the criteria including pregnancy and receiving current or past behavioral therapy for insomnia were excluded in the current study; 4) These participants had not any history of cognitive disorders, neuropsychiatric disorders, or drug abuse and had a suitable family environment for Hemodialysis support.

Of these 590 chronic insomnia patients, 40 chronic insomnia patients (age range: 23-85 years, 36 females, mean insomnia history is 3 years) were collected for the randomized clinical control trials. They were randomly assigned into the test and the control groups (*n* = 20 in each group). Furthermore, we recruited 200 chronic insomnia patients (age from 25-55 years, 128 male/72 female, mean insomnia history is 7 years), we divided these patients, by a pseudo-randomized way, into the test and control groups, via combining the PSQI and the Polysomnography (PSG) to explore the sleep anatomy change underlying the insomnia symptom recovery from 4-week MIR modulation. To trace the dynamic change of the improvement of sleep symptoms coming from MIR biological effect, we recruited 350 chronic insomnia patients for follow-up study (age range: 24-83 years, 168 males) lasting for 8 weeks.

This study was approved by the Ethics Committee of the School of Psychology, South China Normal University and was conducted according to the Declaration of Helsinki. We obtained written informed consent from each participant before the experiment.

### Experimental design

To explore the MIR-induced biological effect, we monitored the blood flow and the blood cell properties using the laser speckle contrast imaging and the microscope with a photographic device. Further more, we explored the modulation effect of MIR on the blood properties in the patients with renal failure in order to track the follow-up effect of the MIR-derived biological effect on human body. To trace the dynamic change of the improvement of sleep symptoms coming from the MIR biological effect, the sleep change from 4-week MIR (1.5 hours each day) was measured using the Pittsburgh Sleep Quality Index (PSQI) and Polysomnography (PSG). Finally, we did a follow-up study by recruiting another 350 chronic insomnia patients (age range: 24-83 years, 168 males, mean insomnia history is over 1 month) to trace the dynamic change of the improvement of sleep symptoms coming from the MIR biological effect within 8 weeks.

### Microscope imaging

We utilized phoenix microscope (PH50) with a photographic device (MC-D500U(C)) on 20 healthy subjects (age range: 22-57, 11 males) to observe the RBC properties.

### Laser speckle contrast imaging

Laser Speckle Contrast Imaging (LSCI) technique which allowed a record of up to 100 images per second was applied to non-invasively monitor the peripheral micro circulatory blood flow in the left hand. Previous studies have shown that LSCI gives very good reproducibility with an excellent spatial and temporal resolutions.(M Roustit, Millet, Blaise, Dufournet, & Cracowski, 2010) LSCI was not only used to estimate the endothelial and neurovascular function, but also as an innovative tool to quantify micro vascular response to clinical treatment.(Matthieu Roustit & Cracowski, 2013) By using the LSCI, we derived the skin microvascular perfusion from speckle contrast analysis with colors ranging from blue (low perfusion) to red (high perfusion), to further provide a perfusion index proportional to skin blood flow. The LSCI was measured in a laboratory with the temperature of 22 ℃.

### Routine Blood Test

The blood cell indices were calculated using a hematology analyzer, including the White Blood Cell Count (WBC), Monocyte percent (MO%), Absolute monocyte count (MONO), Red Blood Cell Count (RBC), RDW, MCHC, Mean Cell Volume (MCV), MCH, Hematocrit (Hct), Lymphocyte percent (LY%), Lymphocyte (LY), BA%, BA, Eosinophil percent (EO%), Eosinophil (EO), Neutrophil percent (NE%), Neutrophil (NE), Hemoglobin Concentration (Hgb), Platelet Count (Plt), Platelet Distribution Width (PDW), Mean Platelet Volume (MPV), and Prothrombin Consumption Time (Pct).

### Concentration of HCY

The HCY was measured by high performance liquid chromatography (HPLC) method with fluorimetric detection based on the Bayer ADVIA Centaur HCY assay. The Bayer ADVIA Centaur HCY assay is a direct chemiluminescent technology based on competitive inhibition. All reduced and bound HCY present in 20 Al of serum or plasma sample is reduced to free HCY. DL-Dithiothreitol (DTT)was used to reduce 20 Al reduced-bound HCY in serum or plasma samples to free HCY with a S-adenosyl-L-homocysteine hydrolase (SAH) in the presence of adenosine (Adn) as substrate, and SAH was used to convert all free HCY enzymes into SAH. A limited number of anti-SAH monoclonal antibody labeled by paramagnetic particles (PMP) coated with biocatalytic SAH-binding avidin produced by patient samples and HCY were used as chemiluminescent tracers. And after the immunization, nine SAHS concentrations ranging from 0 to 65 Amol/l prepared in the protein buffer matrix were magnetic separated and washed to generate standard curves. Multiple repetitions of these criteria are used to define an average standard curve (Master curve). Lot specific master curve information (RLU and concentration) is only provided in the final toolkit. In order to adapt to any instrument to instrument and point-to-point changes, the test includes two calibration devices of specified concentration, and tested at 28-day intervals were conducted to re-calibrate. Control daily operation to ensure that the system is analyzed and calibrated.

### Quantification of T cells

Peripheral blood samples were used for quantification of T cells absolute counts and percentage distribution of lymphocyte subpopulations by flow cytometry. CD3^+^, CD4^+^ and CD8^+^ T lymphocytes were measured using the BD FACSCanto II. Firstly, a special absolute counter was taken with adding 10 *ul* antibody. Secondly, 50 *ul* anticoagulant was added by reverse sampling method that was mixed well in order to avoid encountering globules. Incubation at room temperature and away from light for 30 minutes, followed adding 450 *ul* hemolysin incubation at room temperature and away from light for 5 minutes (CD3-FITC/CD8-PE/CD45-PerCP/CD4-APC). Finally, T cells were tested by FACSCanto.

### Pittsburgh Sleep Quality Index

The Pittsburgh sleep quality index (PSQI) (Buysse, Reynolds III, Monk, Berman, & Kupfer, 1989) was used to measure sleep quality at the time of the clinical interview. The Chinese version of the PSQI scale was used. All participants were instructed to fill the PSQI by a simple demonstration and completed the forms. PSQI scores of the participants was carried out by an experienced sleep researcher blinded to the experimental design, and the global PSQI scores were used to evaluate sleep and sleep characteristics.

### Polysomnography

Polysomnography (PSG) technique with an 18-channel polysomnographic recording system (model 78, Grass Instruments, Quincy, MA) was employed to record the physiologic parameters of sleep. The sleep performance was monitored individually using PSG overnight in a sleep laboratory (a controlled setting under the continued supervision of a sleep technician) during which participants were encouraged to sleep at their own will. The detailed PSG procedures were keeping corresponding to a previous study. (Young et al., 1993) For the present investigation, the following sleep parameters were obtained from the PSG: the total sleep time (TST) in hours; sleep onset latency (SOL), time from lights out to the first epoch of EEG-evidenced sleep in minutes; wake time after sleep onset (WASO), time awake between first and last epochs of EEG-assessed sleep in minutes; sleep efficiency, percent of time in bed spent asleep; and percentage of TST in N1, N2, and N3 stages and rapid eye movement (REM) sleep.

### Five-point Likert-type scale

In order to track the dynamic change of the sleep behavior performance flowing with the bracelet wearing, subjects rated their sleep quality using a 5-point Likert-type scale from 1 (no symptom) to 5 (very severe symptom).(Goldsmith & Levin, 1993; Reyner & Horne, 1995)

### Statistical analysis

All statistical analysis was performed using SPSS Statistics (Version 21.0). According to the distribution of the variables, we used the Analysis of variance (ANOVA, the Bonferroni post hoc Test) and the Student *t* test or the Kruskal–Wallis test and Mann– Whitney *U* test to compare between groups. Paired samples *t*-tests was used to assess differences over time within a group, and a significant level was defined when *P* < 0.05.

## Acknowledgments

We would like to thank all the participants for their involvement.

## Author contributions statement

D.Z. designed the research plan. Y.S., G.X., and C.W. performed all experiments. Y.S. and G.X. performed all the data analysis, Y.S. prepared all the materials, D.Z. wrote the manuscript and supervised all the experiments.

## Additional Information

**Data and materials availability:** All data needed to evaluate the paper are present in the paper and/or the Supplementary Materials. Additional data related to this paper may be requested from the corresponding author.

**Competing interests:** The authors declare that they have no competing interests.

